# Deep Tissue Second-Harmonics of Collagen Fibers in a Transparent Rat Heart from a Myocardial Infarction Model

**DOI:** 10.1101/2025.01.12.631421

**Authors:** Makoto Matsuyama

## Abstract

The combination of tissue-clearing techniques with light-sheet microscopy has enabled detailed visualization of histological changes from the micrometer to millimeter scale, deepening understanding of various disease processes. However, these protocols are not fully optimized for animal species beyond mice or for organs outside the brain. Additionally, the lack of suitable fluorescent probes for target molecules limits their broader application.

In this study, we present a protocol for whole-organ clearing of rat hearts in a myocardial infarction model, achieving complete transparency and enabling label-free imaging of collagen fibers in the myocardial wall up to a depth of ∼5 mm using Second Harmonic Generation (SHG) microscopy. For the first time, we successfully compared collagen fiber orientations between infarcted and healthy myocardial regions. Our approach facilitates high-resolution tissue remodeling analysis in cardiovascular research without the need for antibody staining, demonstrating that tissue-clearing techniques are feasible even in animal species with limited available antibodies.

**Summary statement:** We present a novel method for visualizing collagen fibers deep within the heart using a combination of tissue clearing and advanced microscopy, providing valuable data for cardiac research.

## Introduction

Since the introduction of ultramicroscopy in 2007 [1]—which combined mouse brain-clearing methods with light-sheet microscopy—tissue-clearing technology has rapidly advanced, especially for mouse models. Over the past decade, compounds for delipidation, decolorization, demineralization, and refractive index adjustment have enabled faster and more refined clearing techniques [2–26], allowing entire mouse organs to be rendered transparent postmortem and facilitating single-cell level observations [8,10,11,16,17,23,27–32]. However, protocols optimized for mice often do not directly translate to other species, especially larger animals such as rats.

Rats are frequently used in disease models due to their larger organs and physiological similarities to humans, making them ideal for studies in cardiovascular research [33–35]. Their hearts, for instance, are several times larger than those of mice, posing specific challenges for tissue clearing due to the extended time required for clearing large organs. Additionally, differences in biochemical composition between organs, such as the lipid-rich mouse brain versus hemoglobin- and myoglobin-rich rat hearts and kidneys [36,37], add complexity to the clearing process. Clearing protocols developed for mouse brains are typically inadequate for these larger, more complex rat organs without substantial modifications.

Another limitation in rat models is the shortage of genetic tools and specific fluorescent probes that facilitate high-resolution imaging. Unlike mice, where genetically modified strains expressing fluorescent proteins in targeted cells are readily available, fewer such models exist for rats [38, 39], making antibody staining essential. However, commercially available antibodies validated for use in cleared rat tissues are nonexistent, further complicating the imaging process and limiting specific molecular visualization in these models.

In this study, we present an optimized protocol for whole-organ clearing of rat hearts in a myocardial infarction model, achieving complete transparency. Using label-free Second Harmonic Generation (SHG) microscopy, we enabled high-resolution imaging of collagen fibers within the myocardial wall up to a depth of approximately 5 mm without the need for antibody staining. Additionally, we demonstrated, for the first time, the ability to visualize blood vessels, including newly formed vessels at the infarction site, using only nuclear staining. This approach provides a clear, antibody-free view of vascular remodeling in injured cardiac tissue, yielding new insights into the structural changes associated with infarction.

## Materials and methods

### Animal Experiment

All animal experiments were conducted following the Tokyo Women’s Medical University Animal Experiment Committee guidelines (Approval No. AE19-156, AE20-105, AE21-107, AE22-027, and AE23-124). Sprague-Dawley (SD) and F344/NJcl-rnu/rnu rats (8-10 weeks old) were used to model myocardial infarction (MI). Rats were anesthetized with isoflurane and mechanically ventilated. A left thoracotomy was performed to expose the heart, and the left anterior descending artery was ligated to induce ischemia. Post-surgery, animals were monitored daily. Euthanasia was performed via exsanguination under deep anesthesia one month post-MI. If any rat exhibited a >20% decrease in body weight, it was humanely euthanized as per ethical guidelines. All animals were bred under standard conditions (12-hour light/dark cycle, free access to food and water).

In addition, a fixed marmoset brain previously preserved in 4% paraformaldehyde (PFA) was used for histological analysis.

### Tissue Fixation and Clearing Protocols

Chemical fomula of each reagents followed the CUBIC-Histovision original paper [40]. In this report, three distinct protocols were utilized for tissue clearing:

Protocol 1 (Fig. 1):

1. Delipidation: Excised hearts were washed with PBS containing heparin and fixed in 4% neutral-buffered formalin at 4°C for 3 days. Hearts were washed in PBS (3 × 3 hours) and immersed in 0.5x CUBIC-L overnight at room temperature. Clearing proceeded in 1x CUBIC-L at 45°C for 3 months with bi-weekly solution changes.
2. Nuclear Staining: Cleared hearts were stained with SYTOX-Green (2 μM) or propidium iodide (10 µg/ml) diluted in ScaleCUBIC-1A with 500 mM NaCl for 1 week at 37°C, followed by washing in 10 mM HEPES (pH 7.5) (3 × 3 hours at 25°C).
3. RI Matching: Samples were immersed in 0.5x CUBIC-R+ for 24 hours, then in 1x CUBIC-R+ for 7 days at 25°C.

**Figure 1.**
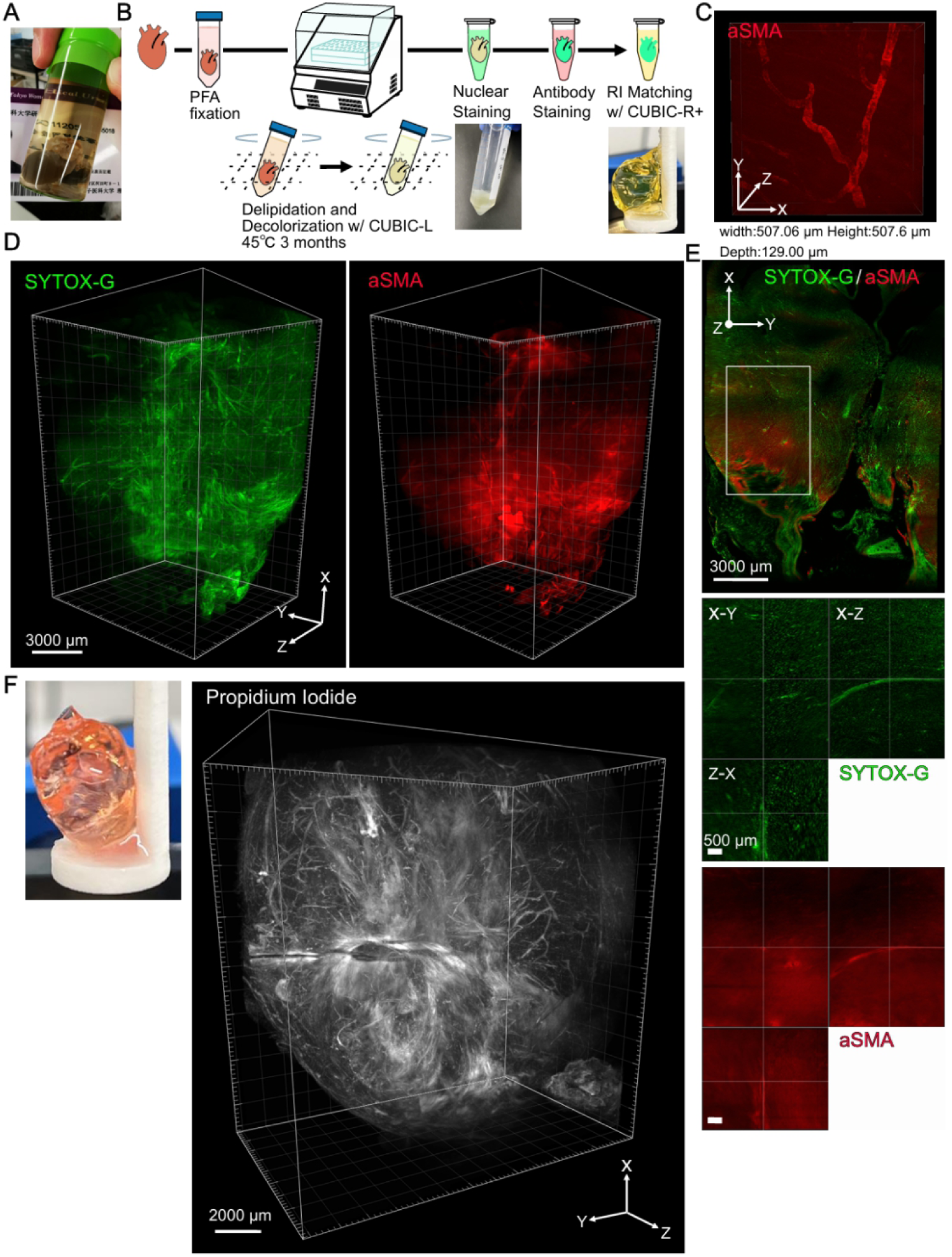
Clearing of whole rat hearts. **A.** Transparent rat hearts processed using a clearing protocol originally optimized for mouse brains. **B.** Workflow for clearing, antibody staining, and nuclear staining of rat hearts. Hearts were immersion-fixed in PFA, subjected to delipidation at 45°C for 3 months, followed by nuclear staining and antibody staining with refractive index matching. **C.** Confocal microscopy image of the vascular structure in a rat heart stained with the vascular marker anti-aSMA antibody. The image was captured 129 µm below the heart surface. **D.** Light-sheet microscopy image of a cleared rat heart stained with SYTOX-Green for nuclei and anti-aSMA antibody for vasculature. The irregular shape of the heart is due to tissue loss during the delipidation process. **E.** Light-sheet microscopy images of the xy-plane of the cleared heart. **F.** Light-sheet microscopy images of a cleared heart stained with propidium iodide. In the myocardial infarction model, the shadow of the ligature used to occlude the vessel is visible, along with vasculature across the heart.

Protocol 2 (Fig. 2):

1. Delipidation: Rats were injected intraperitoneally with 2% Evans Blue in saline (4 mL/kg body weight) 24 hours before euthanasia. Hearts were fixed in 4% formalin at 4°C for 3 days, placed in pyramid tea bags, and cleared in 0.5x CUBIC-L at room temperature overnight. Subsequently, they were cleared in 1x CUBIC-L at 37°C for 3 months with bi-weekly solution changes.
2. Nuclear Staining: Hearts were stained with SYTOX-Green or propidium iodide in ScaleCUBIC-1A with 500 mM NaCl for 3 days at 37°C, followed by washing in 10 mM HEPES (3 × 3 hours at 25°C).
3. RI Matching: Hearts were washed in PBS (1 day at room temperature) and immersed in 0.5x CUBIC-R+ for 24 hours, then in 1x CUBIC-R+ for 3 days at 37°C.

**Figure 2.**
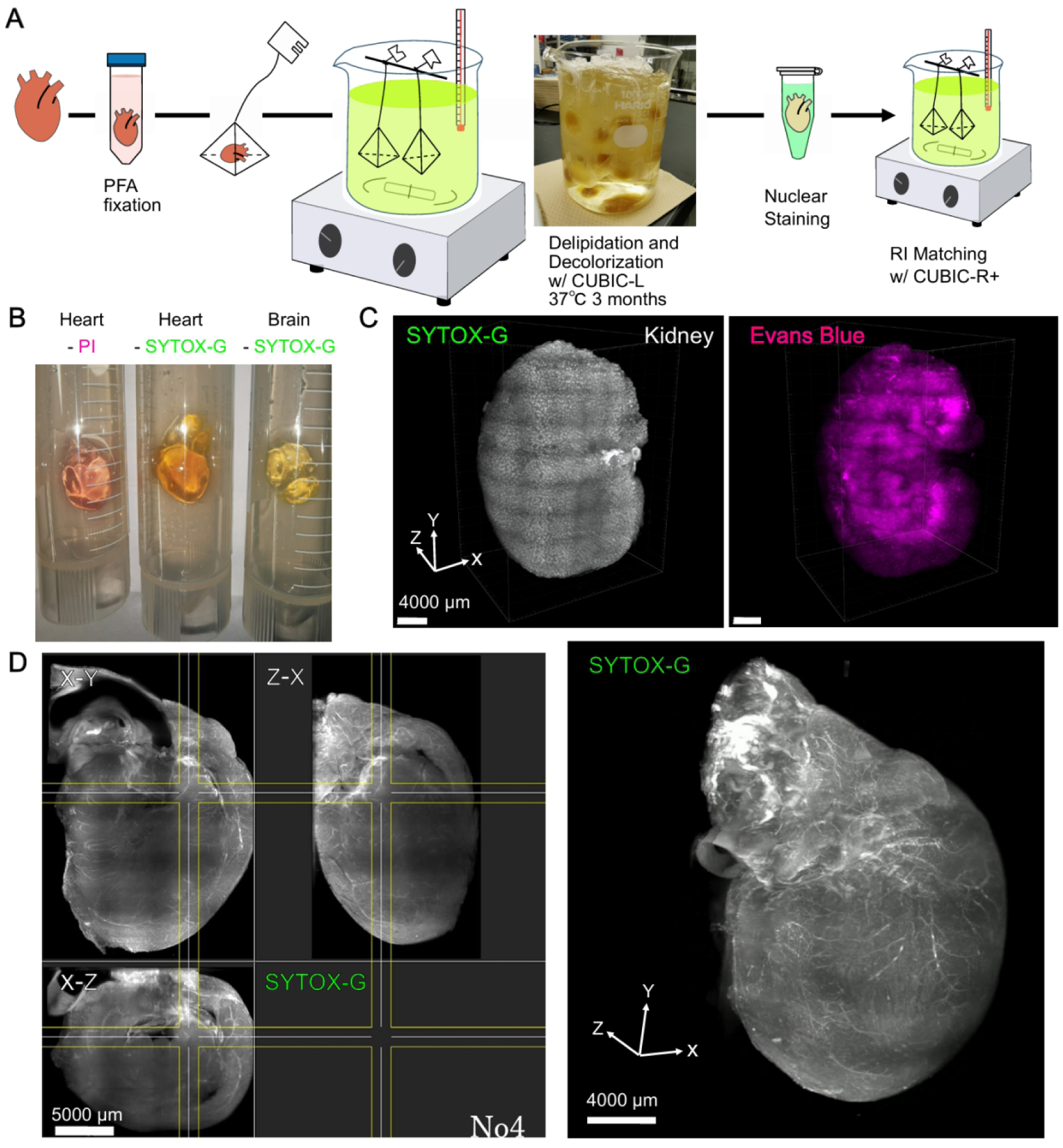
Multi-sample processing using tea bags. **A.** Schematic of the optimized clearing protocol using tea bags for multi-sample processing, including nuclear staining. Rat hearts, kidneys, livers, and brains were immersion-fixed in PFA and subjected to delipidation at 37°C for 3 months, followed by nuclear staining in 1.5 mL tubes and refractive index matching. All other processes were conducted with samples enclosed in tea bags. **B.** Light-sheet microscopy images of rat hearts stained with either SYTOX-G or propidium iodide, and rat brains stained with SYTOX-G using this protocol. **C.** Light-sheet microscopy image of a rat kidney stained with SYTOX-G after pre-labeling blood vessels with Evans Blue. **D.** Light-sheet microscopy image of a whole rat heart stained with SYTOX-Green.

Protocol 3 (Fig. 3):

1. Delipidation: One month after MI induction, perfusion fixation with 4% PFA was performed, followed by perfusion with CUBIC-P solution. Hearts were placed in tea bags and first immersed in 0.5x CUBIC-L at room temperature overnight, then transferred to 1x CUBIC-L in tea bags for 2 months at 37°C, followed by an additional month at 45°C.
2. Nuclear Staining: Hearts were stained with SYTOX Green or propidium iodide in ScaleCUBIC-1A with 500 mM NaCl at 32°C for 3 days, followed by washing as per Protocol 2.
3. RI Matching: As per Protocol 2.

**Figure 3.**
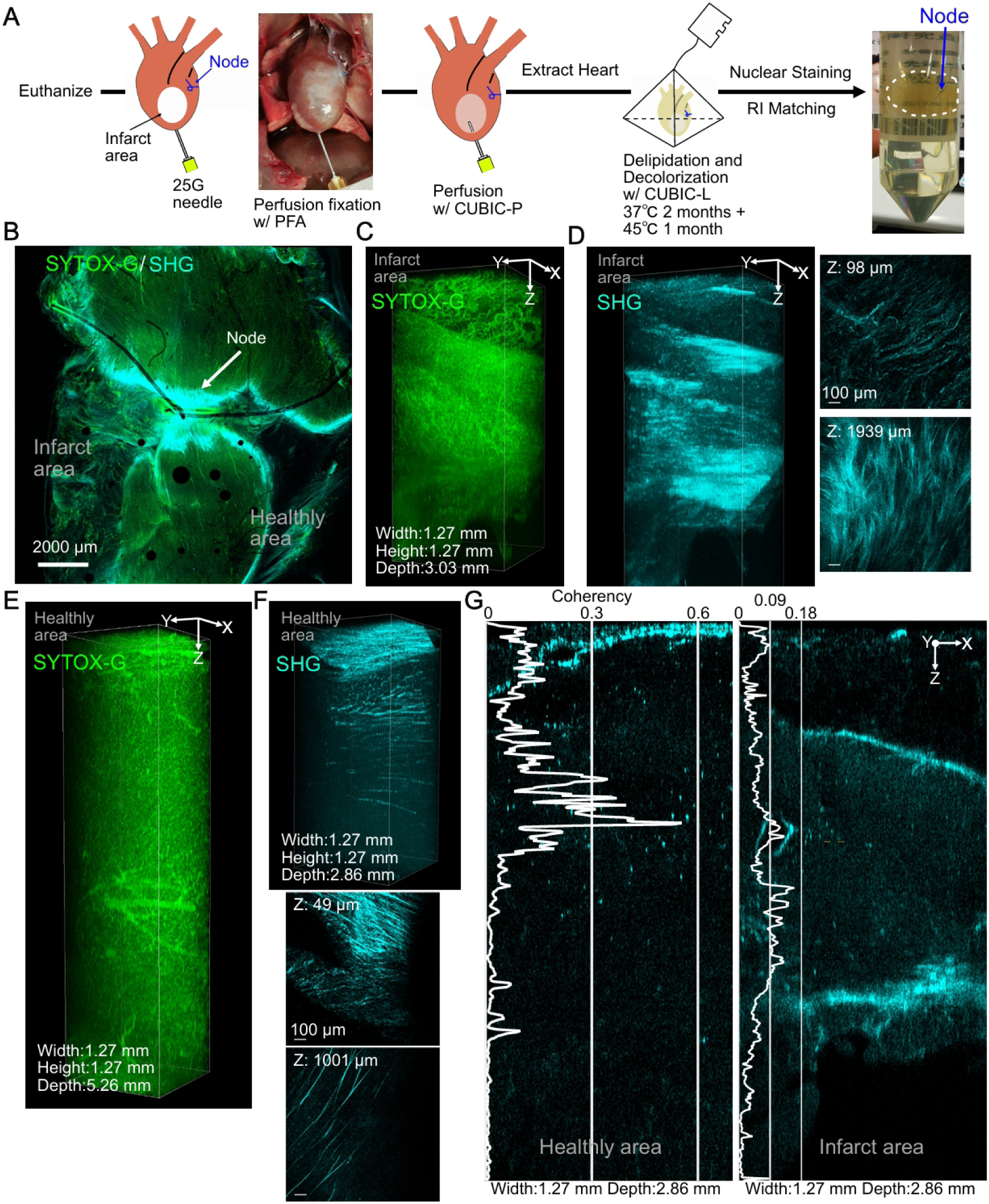
Optimization of perfusion-delipidation and tea bag processing under varied temperature conditions. **A.** Schematic of the optimized clearing protocol for rat hearts. Hearts were perfusion-fixed with PFA and clearing reagents via a 25G needle inserted into the apex. Samples were then processed in tea bags with delipidation at 37°C for 2 months and 45°C for 1 month. **B.** Multi-photon microscopy images of cleared rat hearts stained with SYTOX-Green and imaged for second harmonic generation (SHG). The infarcted region (lower left) downstream of the ligated vessel showed structural replacement by worm-like formations, distinct from the healthy myocardium (lower right). **C.** Nuclear staining image of the infarcted region showing worm-like structures on the surface and myocardial thinning to less than 3 mm. **D.** SHG image of the infarcted region. Short collagen fibers aligned along worm-like structures at 98 µm depth, while densely packed short collagen fibers were observed at 1939 µm depth. **E.** Nuclear staining image of the healthy myocardium showing myocardial thickness exceeding 5 mm, with vasculature evident at both the surface and a depth of 3 mm. **F.** SHG image of the healthy myocardium. Thick, short collagen fibers were observed at a depth of 49 µm, while sparse, long collagen fibers were visible at 1001 µm. **G.** SHG images and collagen fiber coherence in healthy (left) and infarcted (right) regions.

**Figure 4.**
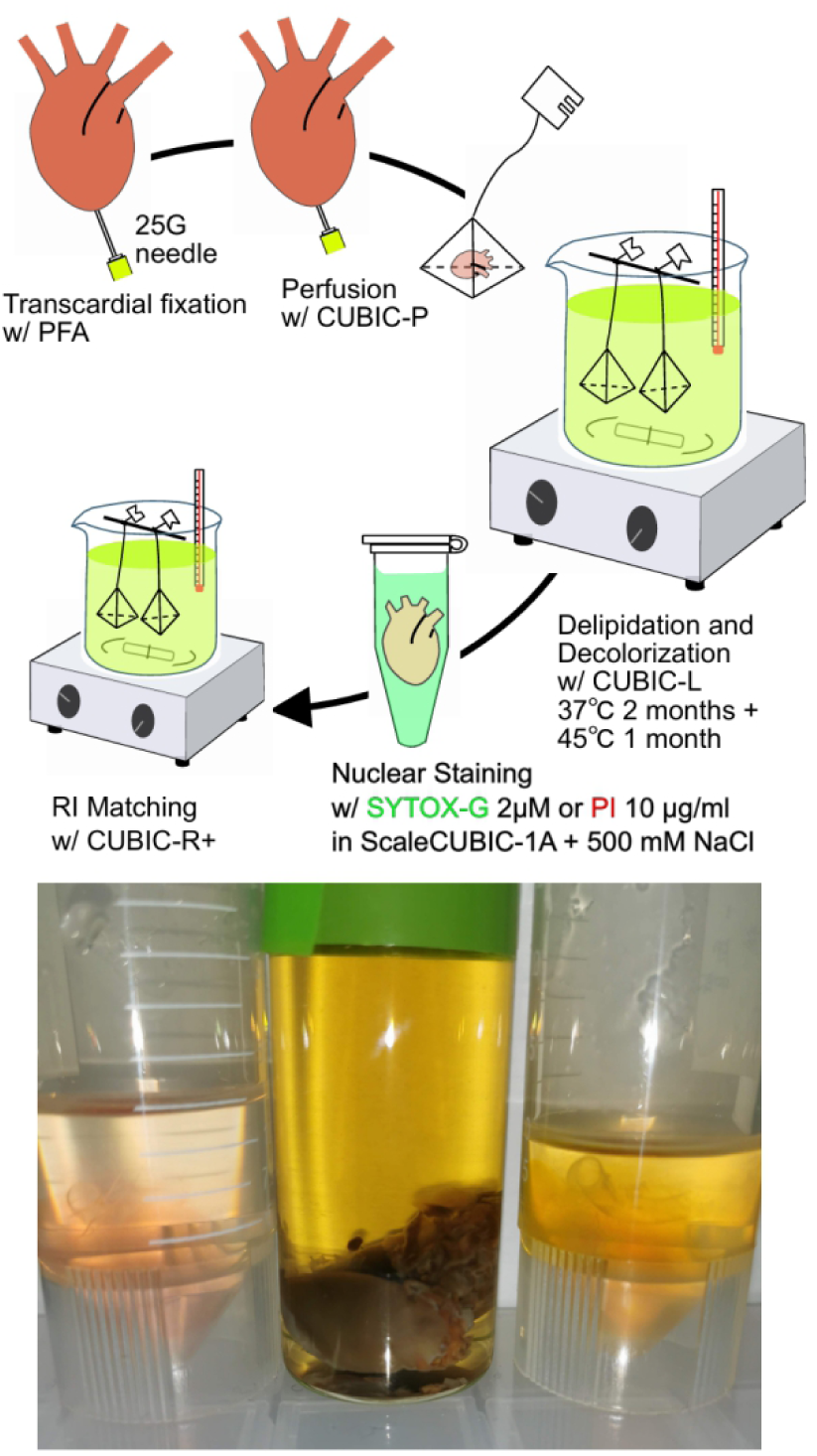
Optimized clearing protocol for whole rat hearts. The optimized protocol employs perfusion-delipidation to improve tissue transparency, physical protection of samples in tea bags during processing, and enhanced multi-sample handling. Delipidation was performed at 37°C for 2 months and 45°C for 1 month to achieve maximum transparency, followed by nuclear staining (PI or SYTOX-G) for 3 days. Images show comparisons of rat hearts processed with the mouse brain protocol (center) and the optimized protocol (left and right). PI-stained heart (left) and SYTOX-G-stained heart (right) demonstrate superior transparency and staining with the optimized protocol.

### Microscopy Setup

Luxendo MuVi SPIM CS (Fig. 1):

- Excitation: 488 nm for SYTOX Green, 561 nm for propidium iodide.
- Detection: Emission filters 525/50 nm for SYTOX Green, 620/60 nm for propidium iodide.
- Objectives: 2×/0.5 NA water immersion objectives.
- Imaging volume: Z-stack with a step size of 2 μm.

LCS SPIM (Fig. 2):

- Excitation: 488 nm (SYTOX Green), 561 nm (propidium iodide).
- Detection: As described for MuVi SPIM.
- Objectives: 4×/0.3 NA water immersion.

Nikon AXR MP multiphoton microscope (Fig. 3):

- Excitation: 920 nm (SHG imaging of collagen).
- Detection: Emission at 450/50 nm for SHG signals.
- Objectives: 25×/1.1 NA water immersion.
- Imaging volume: Z-stack with a step size of 1 μm.

Software: Image analysis was conducted using Imaris 9.9.0, Luxendo Image Processor, and Fiji (ImageJ) with specific plugins, orientationJ, for multiphoton microscopy.

## Results

### Vascular Staining of Rat Organs Using Antibody Immunostaining

Initial attempts to clear rat hearts using a protocol optimized for mouse brain tissue [40] did not yield sufficient results, as the tissue clearing remained incomplete (Fig. 1A). We therefore employed a more aggressive protocol combined with anti-αSMA antibody staining to visualize the vasculature in myocardial infarction model rat hearts (Fig. 1B). Unlike the conventional perfusion fixation commonly used in mouse brain studies, which involves cardiac puncture and could damage the heart tissue, we opted for immersion fixation in paraformaldehyde (PFA) following organ extraction. The tissue clearing process using CUBIC-L solution was extended to three months at 45°C, compared to the standard 3-4 days at 37°C used for mouse brain tissue.

Multiphoton microscopy revealed αSMA-positive vessels in the cleared cardiac surface tissue (Fig. 1C and 1D). While vascular structures were successfully visualized at depths exceeding 3 mm, there was considerable background fluorescence (Fig. 1E). Moreover, the extended CUBIC-L treatment at 45°C resulted in significant tissue deterioration in two out of five samples. Even in successfully imaged samples, mechanical damage was evident where the heart tissue had repeatedly contacted the tube walls during agitation (Fig. 1D and 1F).

Notably, nuclear staining with SYTOX-G and Propidium Iodide (PI) not only labeled cardiomyocyte nuclei but also provided exceptional visualization of the vasculature with high signal intensity (Fig. 1D and 1F) and maintained low background fluorescence even in deep tissue regions (Fig. 1E).

### Evans Blue Staining for Vascular Visualization in Rat Organs

We next explored Evans Blue staining as an alternative vascular labeling method, utilizing its property of binding to blood albumin and remaining within vessels. This approach was tested in healthy animals to avoid potential dye leakage in infarcted regions. To minimize mechanical damage during long-term agitation, we developed an improved clearing method using pyramid-shaped tea bags (Fig. 2A). This setup, comprising a large beaker filled with CUBIC-L solution and suspended tea bags on a hot stirrer, facilitated simultaneous processing of multiple samples. We applied this method to clear and nuclear-stain rat brain and kidney tissues in addition to heart tissue (Fig. 2B and 2C), reducing the temperature to 37°C for a three-month period.

While the clearing process was successful, the Evans Blue dye was largely eliminated during the three-month clearing period, resulting in only minimal vessel visualization (Fig. 2C). Nuclear staining patterns varied by tissue type: kidney samples predominantly showed glomerular structures rather than strong vascular signals, while heart samples maintained excellent surface tissue integrity, demonstrating the effectiveness of the tea bag method. The nuclear staining alone continued to provide detailed vascular visualization (Fig. 2D). However, compared to Figure 1, deep tissue transparency was reduced with higher background fluorescence, indicating that temperature conditions required further optimization.

### Optimized Protocol and Second Harmonic Generation Imaging of Collagen Fibers

Having established that nuclear staining alone could effectively visualize cardiac vasculature, we focused on maximizing tissue transparency for deep tissue imaging. We modified our protocol to include perfusion fixation with PFA via cardiac apex injection, followed by immediate perfusion with CUBIC-P solution for delipidation and decolorization (Fig. 3A). This approach improved tissue fixation and initiated clearing before organ extraction. The clearing process was adjusted to include two months at 37°C followed by one month at 45°C to preserve tissue integrity.

This optimized protocol yielded excellent results, producing highly transparent organs without tissue degradation. Using multiphoton microscopy with second harmonic generation (SHG), we observed that SYTOX-G nuclear staining revealed a reduction in myocardial thickness from >5mm in healthy regions to <3mm in infarcted areas (Fig. 3C and 3E). The infarct surface displayed numerous worm-like vascular structures absent in healthy tissue (Fig. 3C). SHG imaging achieved approximately 5mm imaging depth (Fig. 3D and 3F, and Supplementary Movie 5 and 6) and revealed distinct collagen fiber patterns between healthy and infarcted regions. Healthy tissue showed aligned, thick collagen fibers at depths of 782-1396μm, while infarcted regions displayed dense accumulations of shorter, thinner fibers at corresponding depths (Fig. 3G).

Our findings demonstrate that nuclear staining alone can effectively reveal vascular structures in rat cardiac tissue, enabling simultaneous analysis of myocardial and vascular structural changes from the same dataset. The optimization of clearing temperature conditions and the transition from tubes to tea bags has enabled processing of multiple samples without physical damage, shifting the analysis paradigm from representative tissue sections to comprehensive whole-organ analysis without limitations on sample numbers. Furthermore, the achieved tissue transparency has enabled deep-tissue SHG imaging, previously limited to thin skin samples, allowing visualization of individual collagen fibers at depths of several millimeters. This protocol not only liberates researchers from the labor-intensive processes of sectioning, staining, imaging, and reconstruction but also provides unprecedented detail in analyzing collagen deposition in damaged tissue regions, surpassing the capabilities of conventional methods.

## Discussion

This study represents the first successful demonstration of tissue clearing in rat cardiac tissue. While limited reports of tissue clearing in rat organs exist [11,24,41], none have achieved the level of transparency required for detailed imaging of the heart. Additionally, we introduce two significant methodological advances: the first use of nuclear staining to effectively visualize vascular structures, and the first application of second harmonic generation (SHG) microscopy for imaging collagen fibers up to ∼5 mm depth in cleared organs.

Over the past decade, tissue clearing techniques have advanced significantly, evolving through several phases: initial conceptual approaches [1–6], innovations to preserve fluorescent proteins during clearing [25,42,43], and the development of antibody staining methods for cleared organs and whole organisms [31,40,44,45]. Despite these achievements, the field still faces critical challenges, particularly the lack of standardized protocols tailored to different tissue types. This absence forces researchers to overcome numerous technical hurdles to adapt clearing techniques for specific experimental needs. These challenges are especially problematic in the context of animal experiments, where tissue analysis often occurs at the final stages using irreplaceable samples. Understandably, researchers are reluctant to risk such valuable specimens on unproven experimental clearing protocols, particularly when established sectioning methods are available and reliable.

Validated clearing and staining protocols specific to different tissues, along with clear demonstrations of their advantages over traditional methods, are essential for wider adoption. Our experience across various organ types reveals significant tissue-specific differences in clearing processes. For example, brain white matter requires longer clearing times than gray matter, particularly in adult marmoset or macaque brains (SFig. 1E). Lung tissue clears faster but exhibits strong autofluorescence (SFig. 1A and SFig. 2). Liver tissue, despite its high lipid content, becomes too fragile for practical handling during CUBIC clearing (SFig. 1B). Heart and kidney tissues achieve excellent transparency and structural integrity but are highly sensitive to temperature during delipidation and require the longest clearing times among the organs examined (SFig. 1A).

Although our three-month protocol may appear lengthy, it offers practical advantages. Using the pyramid-shaped “tea bag” method, we processed 12 rat heart samples with minimal hands-on time—approximately one hour per week for buffer exchanges. In contrast, traditional sectioning methods for equivalent analyses demand continuous attention for 4–6 months, hindering parallel experimental work. The pyramid design enhances buffer circulation, minimizes physical stress on tissues, and allows simultaneous processing of multiple samples (e.g., 17 hearts, 2 brains, and 2 kidneys in a single 1L beaker) (Fig. 2A). Additionally, this setup eliminates the need for direct tissue handling with forceps, reducing the risk of sample damage.

One unexpected finding was the utility of nuclear staining for vascular visualization in cardiac tissue. When paired with ScaleCUBIC-1A containing 500 mM NaCl, nuclear stains achieved rapid, uniform penetration of large organs with minimal background fluorescence. While vascular visualization is not exclusive to cardiac tissue—kidney samples revealed glomerular structures, and brain samples displayed vessels and neurons (SFig. 1C)—the method’s effectiveness in heart tissue likely stems from the sparse distribution of cardiomyocyte nuclei contrasted with the densely packed endothelial nuclei.

Combining tissue clearing with SHG microscopy enabled imaging of collagen structures at unprecedented depths (∼5 mm), far surpassing the previous limit of 200–300 µm in skin [46]. This approach uncovered distinct collagen fiber patterns in healthy and infarcted ventricular walls, from the epicardium to the endocardium (Fig. 3G). Healthy tissue exhibited ordered, elongated fibers, whereas infarcted regions showed shorter, disorganized fibers. These findings open new avenues for studying tissue fibrosis and remodeling in various pathological conditions.

Despite these advances, challenges remain for broader adoption of tissue clearing techniques:

1. Protocol Duration: Although the three-month clearing process is justified by minimal labor and high-quality results, shorter protocols would improve practicality. Current reagents optimized for lipid-rich brain tissue may not be ideal for organs like the heart and kidney, suggesting the need for tissue-specific clearing agents to reduce processing times [47].
2. Autofluorescence: Deep tissue antibody staining is limited by high background fluorescence (Fig. 1). While photobleaching [48,49] and chemical treatments [50,51] show potential, further optimization is needed, particularly for highly autofluorescent tissues such as the lung (SFig. 2).
3. Fluorescent Probe Development: The success of phalloidin, a middle-sized cyclic peptide, highlights opportunities for developing new tissue-specific probes. While phalloidin’s utility in cardiac tissue is limited by widespread actin staining (SFig. 1D), its rapid penetration and high signal-to-noise ratio in lung tissue, particularly for PDPN and FLT4 double-positive vessels (SFig. 3), underscores the potential of middle-sized peptide probes over conventional heavy-chain antibodies.

This study addresses critical gaps in tissue clearing by providing an optimized protocol for rat cardiac tissue, an efficient multi-sample processing setup, and novel applications in vascular and collagen imaging. These advancements not only broaden the applicability of tissue-clearing techniques but also lay the groundwork for addressing persistent challenges such as autofluorescence reduction and the development of novel fluorescent probes, ultimately enhancing the precision and versatility of cleared tissue imaging in diverse research contexts.

## Acknowledgments

We would like to express our gratitude to Dr. Jun Homma (Tokyo Women’s Medical University and Waseda University Joint Institution for Advanced Biomedical Sciences) for his guidance on the myocardial infarction model preparation. We also thank Luxendo for their support with light sheet microscopy imaging and Nikon for their assistance with multiphoton microscopy imaging.

## Data and resource availability

Imaging data and movies presented in figures are available in a Synapse by Sage Bionetworks (https://doi.org/10.7303/syn64014998).

## Diversity and inclusion statement

Not applicable

---

**Supplementary Figure 1.**
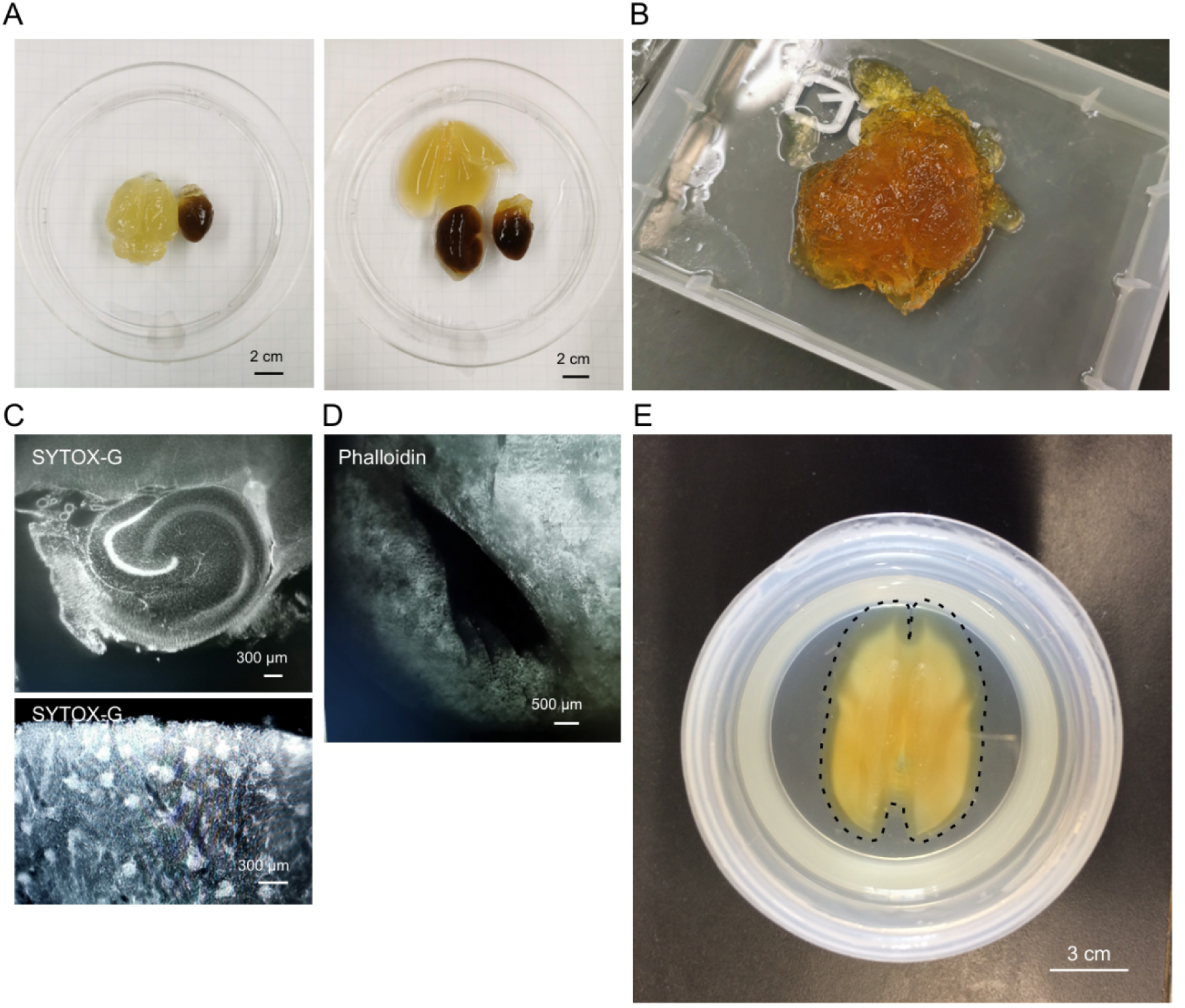
Comparison of tissue transparency during clearing across various rat organs. **A.** Transparency progress of simultaneously processed brain, heart, lung, kidney, and liver tissues. Brains and lungs achieved transparency quickly, whereas kidneys and hearts remained opaque. **B.** Liver tissues became fragile after delipidation, breaking during removal from tea bags despite remaining intact during processing. **C.** Light-sheet microscopy images of cleared rat brains and kidneys stained with SYTOX-Green. Vascular and cellular structures were visible in the brain, and glomeruli were visualized in the kidney. **D.** Phalloidin staining of cleared rat hearts showing F-actin. Although striated structures were discernible, staining appeared diffuse overall. **E.** Transparency of adult marmoset brains processed for 1 week at 37°C. Cortical regions became sufficiently transparent, whereas white matter remained slightly opaque.

**Supplementary Figure 2.**
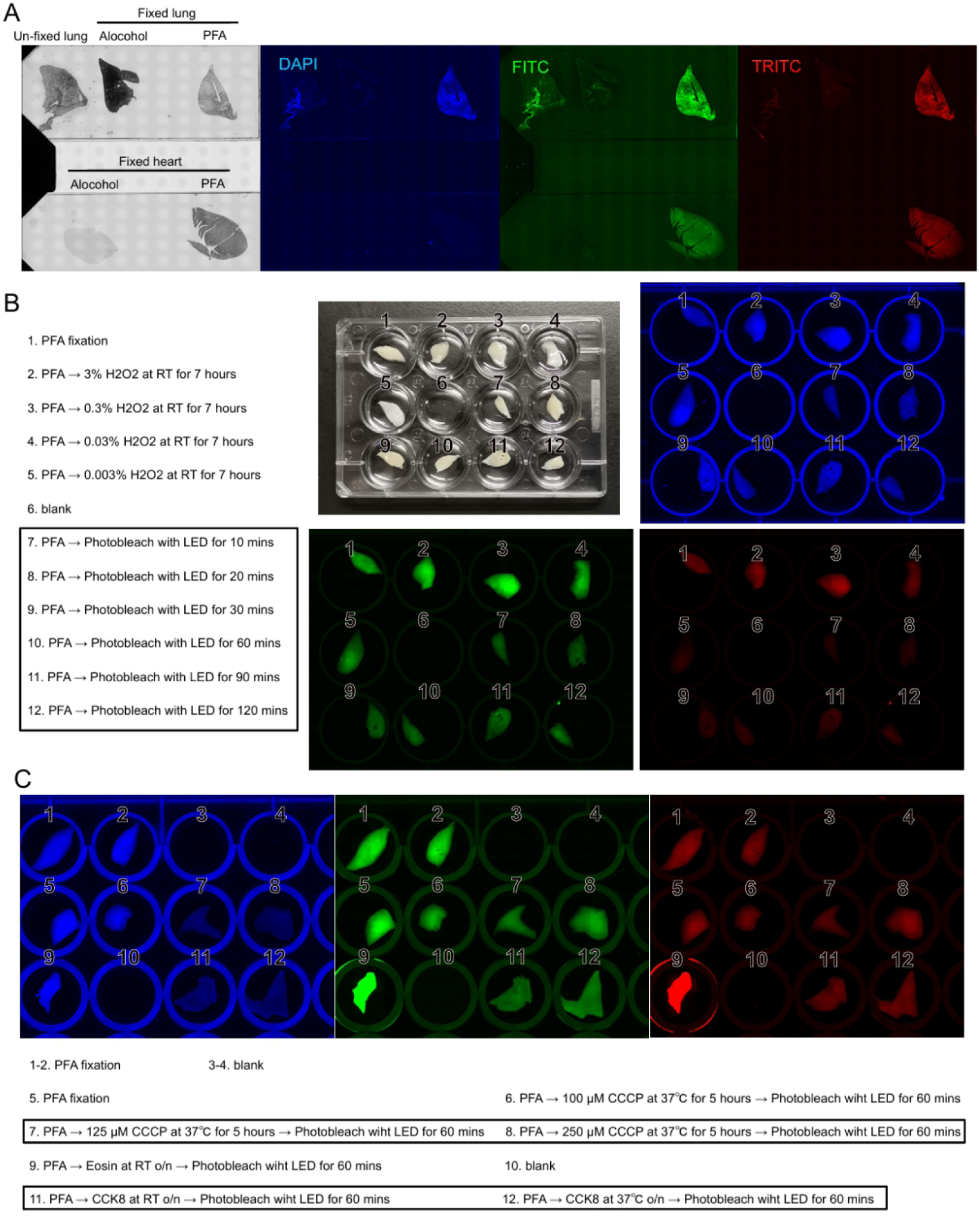
Autofluorescence and quenching strategies for rat lungs. **A.** Autofluorescence of rat lungs and hearts under different fixation methods. PFA fixation resulted in strong autofluorescence across the DAPI, FITC, and TRITC channels, particularly in lungs. This autofluorescence could not be quenched using TrueBlack®, a reagent designed to quench lipofuscin-related autofluorescence. **B.** Attempts to reduce autofluorescence using hydrogen peroxide (0.003–3%) or LED light exposure. Hydrogen peroxide treatment caused the lungs to float, reducing its effectiveness, while LED light exposure successfully diminished autofluorescence in the FITC and TRITC channels but had no effect on the DAPI channel. **C.** Treatment combining CCCP or CCK8, which oxidize NAD(P)H to NAD(P)+, with LED light exposure effectively reduced autofluorescence across all channels under specific conditions. In contrast, the use of Eosin Y, a photoreduction catalyst, followed by LED exposure increased background fluorescence due to the intrinsic fluorescence of Eosin Y itself.

**Supplementary Figure 3.**
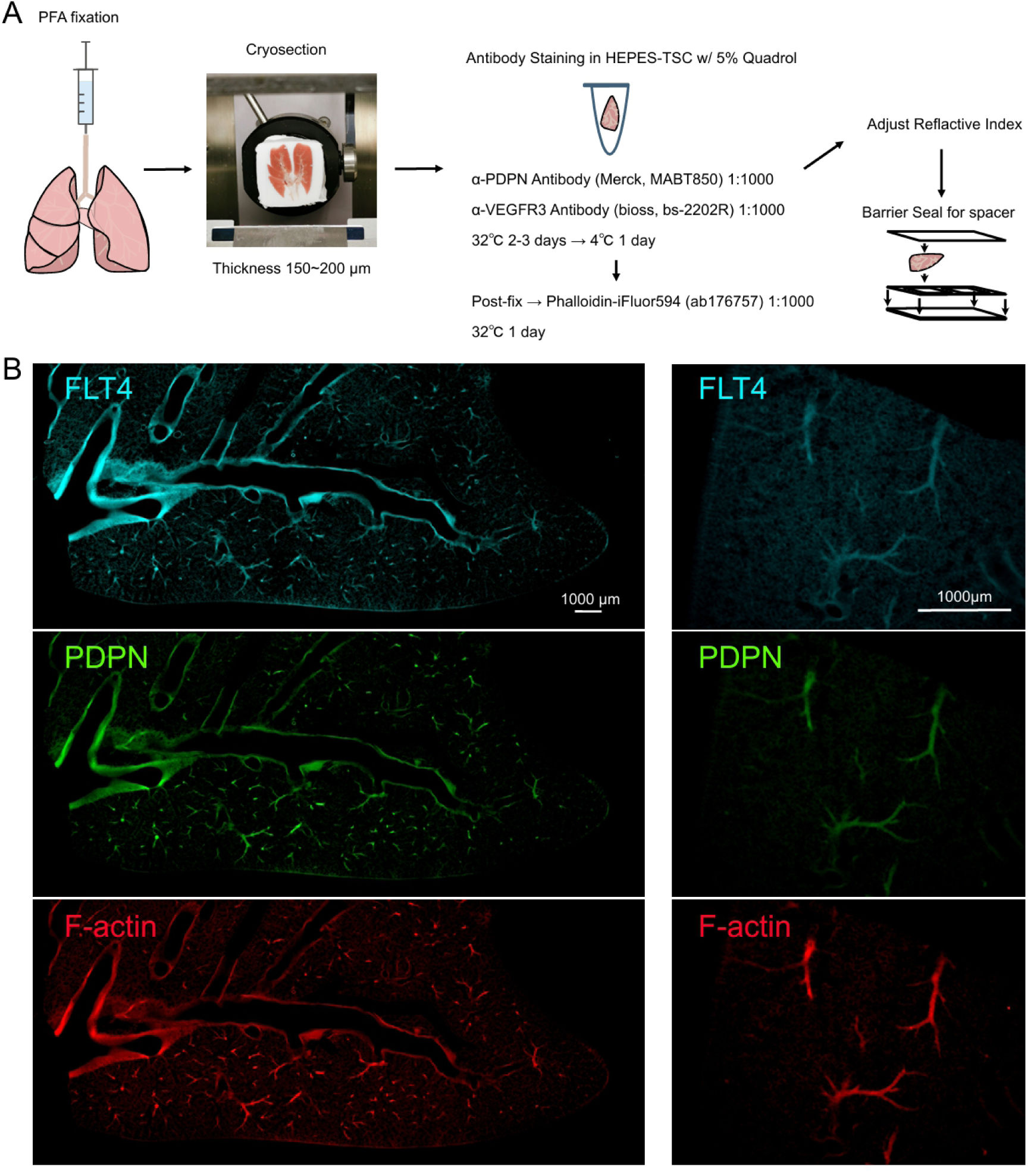
Optimal protocols for vascular visualization in rat lungs. **A.** Whole-lung transparency is insufficient for detailed vessel visualization due to strong autofluorescence. Instead, 150-200 µm sections without delipidation were stained with anti-FLT4, anti-PDPN antibodies, and phalloidin, followed by post-fixation in PFA and mounting with refractive index matching medium. **B.** Multi-channel fluorescence images of 200 µm lung sections stained for FLT4, PDPN, and F-actin. Separate staining of antibodies and phalloidin was essential to avoid granular artifacts.

Supplementary Movie 1

Light-sheet microscopy video of a cleared rat heart stained with SYTOX-Green for nuclei and anti-aSMA antibody for vasculature. Corresponds to Figure 1D.

Supplementary Movie 2

Light-sheet microscopy video of a cleared rat heart stained with PI for nuclei. Corresponds to Figure 1F.

Supplementary Movie 3

Light-sheet microscopy video of a cleared rat kidney stained with SYTOX-Green for nuclei and Evans Blue for vasculature. Corresponds to Figure 2C.

Supplementary Movie 4

Light-sheet microscopy video of a cleared rat heart stained with SYTOX-Green for nuclei and visualizing vasculature. Corresponds to Figure 2D.

Supplementary Movie 5

Multiphoton excitation microscopy video of the infarcted region of a cleared rat heart stained with SYTOX-Green for nuclei. Corresponds to Figure 3C-D.

Supplementary Movie 6

Multiphoton excitation microscopy video of the infarcted region of a cleared rat heart stained with SYTOX-Green for nuclei and visualizing collagen fibers with SHG. Corresponds to Figure 3C-D.

Supplementary Movie 7

Multiphoton excitation microscopy video of the intact region of a cleared rat heart stained with SYTOX-Green for nuclei. Corresponds to Figure 3E-F.

Supplementary Movie 8

Multiphoton excitation microscopy video of the intact region of a cleared rat heart stained with SYTOX-Green for nuclei and visualizing collagen fibers with SHG. Corresponds to Figure 3E-F.

